# *rcsB* enhances drug resistance of *Klebsiella pneumoniae* ATCC43816 by regulating the formation of its capsule

**DOI:** 10.1101/2022.12.24.521839

**Authors:** Huiqing Huang, Tao Wang, Jie Zhu, Shumin Liu, Hong Du

## Abstract

**Objective:** To analyze the composition and drug resistance characteristics of clinical isolates from a tertiary general hospital of Lianyungang in 2021, and to explore the possible regulatory mechanism of drug resistance of *Klebsiella pneumoniae*.

**Methods:** The clinical samples submitted in 2021 were collected for bacterial culture, identification and drug sensitivity test, and the distribution and drug resistance of the isolated pathogens were analyzed. The biological characteristics of highly virulent *Klebsiella pneumoniae* ATCC43816 and its *rcsB* deletion strains were compared by growth curve test, viscosity semi quantitative test, uronic acid quantitative test and drug sensitivity test.

**Results:** A total of 1,077 strains of pathogenic bacteria were isolated, and the top 3 pathogens were *Escherichia coli, Klebsiella pneumoniae, Pseudomonas aeruginosa* (23.96%, 17.46%, 11.23%). *Klebsiella pneumoniae* had high resistance rates to cefazolin, ampicillin/sulbactam and cefuroxime. Compared with the wild strain, the MIC values of imipenem, ampicillin/sulbactam and tegacyclin in Δ*rcsB* was decreased.

**Conclusion:** *Klebsiella pneumoniae* should be the focus of bacterial drug resistance monitoring in order to guide early anti-infection treatment. *rcsB* may indirectly regulate the drug resistance of *Klebsiella pneumoniae* by regulating the amount of capsule formation, which is of great significance for improving the drug resistance status of *Klebsiella pneumoniae*.

## Introduction

With the extensive use of broad-spectrum antibiotics, bacterial resistance has become an increasingly serious public health problem. The summary of the results of bacterial culture in this region and the analysis of drug resistance data are one of the important prevention and control measures. Therefore, we conducted a retrospective analysis of the strain related data of a tertiary general hospital in Lianyungang in 2021. The mechanisms that caused bacterial resistance include the production of hydrolase, the change of drug action structure, the change of bacterial cell membrane permeability, the spread of integrons, *etc*^[1]^. The use of some means to inhibit the expression of corresponding virulence genes may break the situation of multi drug resistant bacteria. *Klebsiella pneumoniae* is one of the common clinical pathogenic bacteria, and ATCC43816 is a highly virulent *Klebsiella pneumoniae*. By comparing the biological characteristics of its capsule regulatory gene *rcsB* defective strain and wild strain, we discussed the possible drug resistance mechanism, hoping to enlighten the control of drug resistance of pathogenic bacteria.

## Materials and methods

### Source of strains

Collect the pathogenic bacteria detected by inpatients in 2021, eliminate duplicate strains, and screen 1,077 strains in total. *Klebsiella pneumoniae* ATCC43816, ATCC43816 Δ*rcsB* comes from the laboratory inventory.

### Test method

Strain identification and drug sensitivity test were conducted according to the National Standard Operating Procedures for Clinical Use (Version 4) for smear, inoculation and separation. The Dade Behring Microscan Autoscan 4 bacteria identification instrument and BDPhoenix-100 automatic bacteria identification instrument were used for identification and drug sensitivity.

### Growth curve of Klebsiella pneumoniae ATCC43816 ΔrcsB

ATCC43816 and Δ*rcsB* were transferred into 20 ml of fresh LB liquid culture medium at a ratio of 1:100 for overnight bacterial solution, placed them in a constant temperature shaking table at 37°C and shaked them at 250 rpm, and 200 μL was taken from them every 1 h. The OD600 of the bacterial solution was measured, and the measured values at each time point within 24 hours are drawn into a growth curve.

### Viscosity semi quantitative test of Klebsiella pneumoniae ATCC43816 ΔrcsB

The OD600 of ATCC43816 and Δ*rcsB* incubated for 8 h were adjusted to 1.0, and then 1,000 g centrifuged for 5 min, and the upper liquid was taken to measure its OD600.

### Quantitative assay of uronic acid in Klebsiella pneumoniae ATCC43816 ΔrcsB

100 μL 1% amphoteric detergent 3-14 and 500 μL overnight bacterial solution of ATCC43816 and Δ*rcsB* were mixed, incubated at 50°C for 30 min, centrifuged at 14,000 rpm for 2 min, and then 250 μL the supernatant was transferred to a new 1.5 ml centrifuge tube, 1 ml of precooled absolute ethanol was added, incubated at 4°C for 30 min to produce sediment, centrifuged at 14,000 rpm for 5 min, discarded the supernatant, left for 3 min, and dissolved the sediment in 200 μL Deionized water. 1.2 ml of sodium tetraborate/concentrated sulfuric acid solution was added to the extract, shaked and mixed, boiled for 5 min, cooled for 5 min, and added 20 μL 3-hydroxydiphenol. After reaction, the absorbance value at 520 nm was measured.

### Statistical treatment

GraphPad Prism 8 was used for data analysis.

## Results

In 2021, 1,077 strains of pathogenic bacteria were detected, including *Escherichia coli, Klebsiella pneumoniae, Pseudomonas aeruginosa, Acinetobacter baumannii, Proteus mirabilis* and *Staphylococcus aureus* (Table 1).

**Table 1.** Composition of main pathogens in 2021.

| Bacterial name | Total number | Proportion (%) |
| --- | --- | --- |
| *Escherichia coli* | 258 | 23.96 |
| *Klebsiella pneumoniae* | 188 | 17.46 |
| *Pseudomonas aeruginosa* | 121 | 11.23 |
| *Acinetobacter baumannii* | 76 | 7.06 |
| *Proteus mirabilis* | 61 | 5.66 |
| *Staphylococcus aureus* | 57 | 5.29 |
| *Staphylococcus epidermidis* | 43 | 3.99 |
| *Enterococcus faecium* | 36 | 3.34 |
| *Enterococcus faecalis* | 34 | 3.16 |
| *Enterobacter cloacae* | 31 | 2.88 |
| *Stenotrophomonas maltophilia* | 21 | 1.95 |
| *Staphylococcus haemolyticus* | 21 | 1.95 |
| *Klebsiella aerogenes* | 20 | 1.86 |
| *Klebsiella oxytoca* | 15 | 1.39 |
| other | 95 | 8.82 |

Resistance of *Klebsiella pneumoniae* to cefazolin, ampicillin/sulbactam, cefuroxime, furantoin, cefotaxime, and ceftriaxone was relatively high. The detection rate of carbapenem resistant *Klebsiella pneumoniae* was 40.0%, and *Klebsiella pneumoniae* was highly sensitive to tetracyclines (Table 2).

**Table 2.** Drug sensitivity analysis of *Klebsiella pneumoniae* (%)

| Antibacterial | R (drug resistance rate)% | I (moderate sensitivity)% | S (sensitivity rate)% |
| --- | --- | --- | --- |
| amikacin | 35.2 | 0.8 | 64.0 |
| ampicillin/sulbactam | 57.3 | 7.3 | 35.4 |
| aztreonam | 54.5 | 1.6 | 43.9 |
| furantoin | 56.0 | 19.0 | 25.0 |
| compound<br>sulfamethoxazole | 38.6 | 0 | 61.4 |
| ciprofloxacin | 53.3 | 2.1 | 44.6 |
| piperacillin/tazobactam | 44.2 | 1.2 | 54.6 |
| gentamicin | 43.9 | 0.8 | 55.3 |
| cefepime | 52.9 | 0.4 | 46.7 |
| ceftriaxone | 55.0 | 0.9 | 44.1 |
| cefotaxime | 55.6 | 0 | 44.4 |
| ceftazidime | 49.3 | 2.4 | 48.3 |
| cefoxitin | 47.1 | 0.9 | 52.0 |
| cefazolin | 57.6 | 0 | 42.4 |
| tobramycin | 39.4 | 2.2 | 58.4 |
| imipenem | 39.4 | 0 | 60.6 |
| levofloxacin | 48.5 | 2.4 | 49.1 |
| cefuroxime | 56.3 | 2.5 | 41.2 |
| meropenem | 40.0 | 0 | 60.0 |
| tigecycline | 10.6 | 0 | 89.4 |

### Growth characteristics of *Klebsiella pneumoniae* Δ*rcsB*

The growth monitoring results of wild strains ATCC43816 and Δ*rcsB* were drawn into a growth curve (Figure 1). There was no significant difference between the growth rate of the deleted strain and the wild strain, indicating that the growth of highly virulent *Klebsiella pneumoniae* ATCC43816 did not change significantly when the *rcsB* gene was missing.

**Figure 1.**
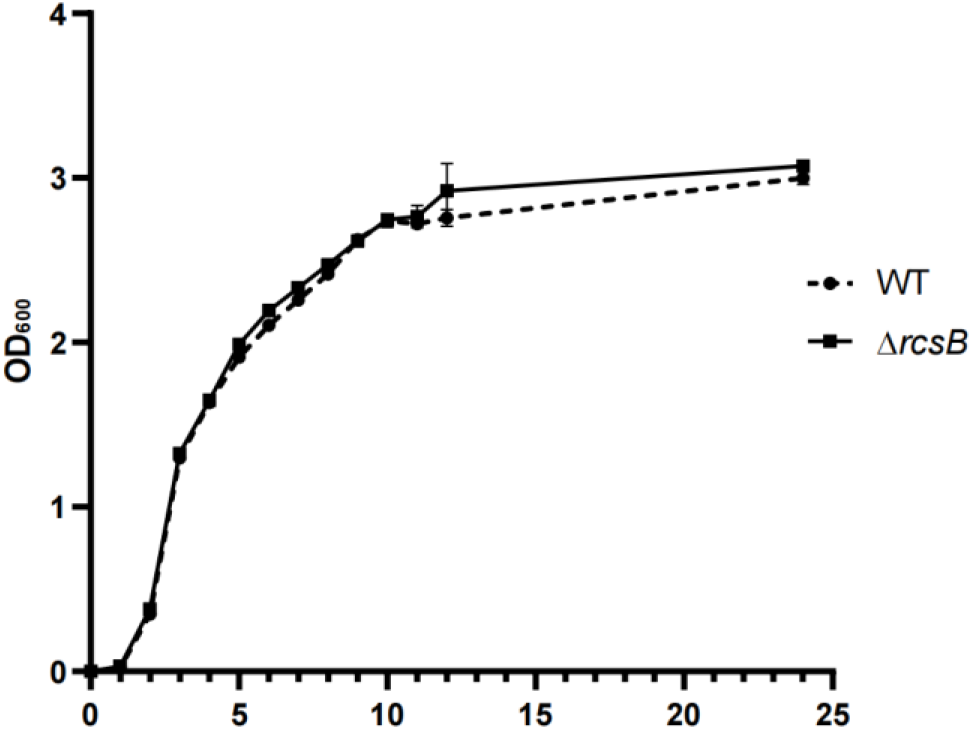
Growth curve of wild strain ATCC43816 and *rcsB* gene deletion strain.

### Viscosity related characteristics of *Klebsiella pneumoniae* ATCC43816 and Δ*rcsB*

The viscosity of each strain was measured by viscosity semi quantitative experiment. After low-speed centrifugation, the suspension of high viscosity strain was more difficult to settle than that of non-high viscosity strain. The viscosity of the strain was quantified by measuring the absorbance value of its supernatant. The results showed that compared with wild strains, the OD600 of *rcsB* gene deletion strains was significantly reduced (Figure 2).

**Figure 2.**
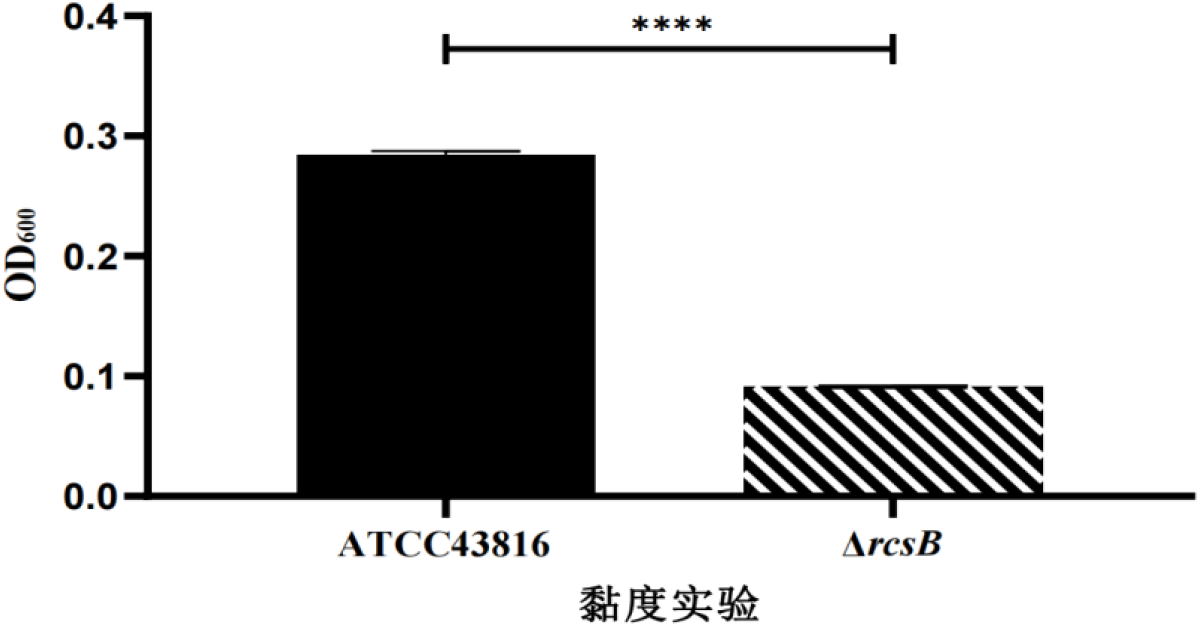
Semi-quantitative results of viscosity of wild strains ATCC43816 and *rcsB* gene deletion strains (**** *p*<0.0001, *t*-test)

### Quantitative results of uronic acid in *Klebsiella pneumoniae* ATCC43816 and Δ*rcsB*

The quantitative results of glucuronic acid showed that the quantitative results of *rcsB* gene deleted strains were significantly lower than those of wild strains, indicating that the deletion of *rcsB* gene led to the decrease of capsule yield of highly virulent *Klebsiella pneumoniae* (Figure 3).

**Figure 3.**
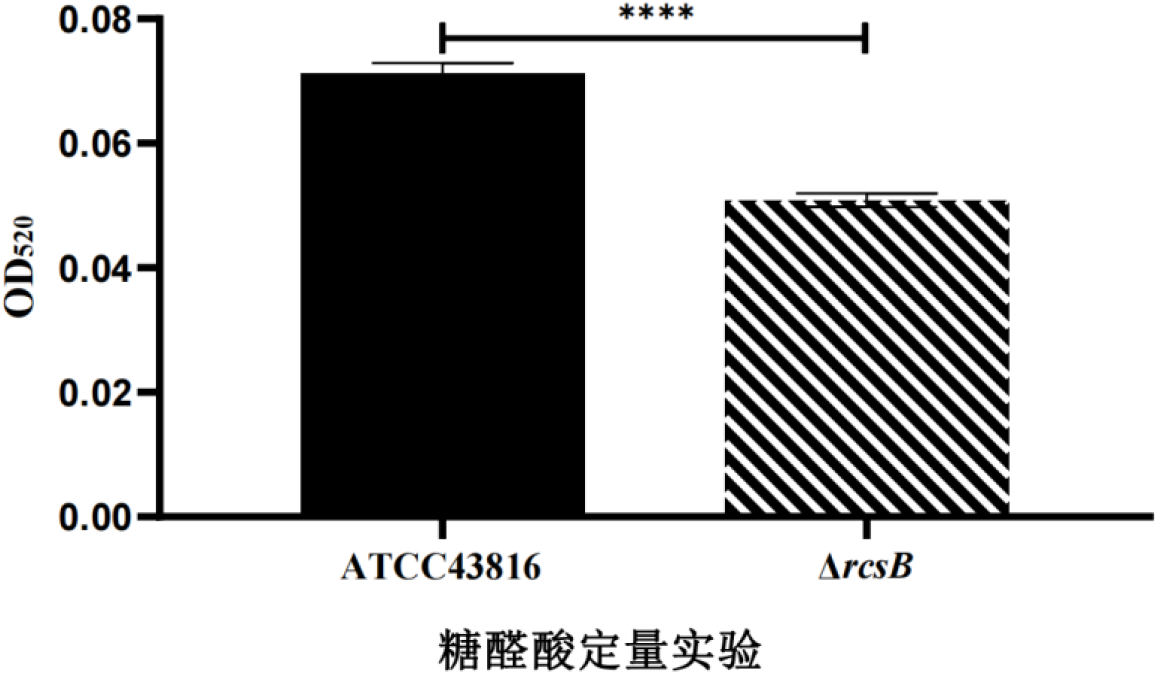
Quantitative results of glucuronic acid of wild strains ATCC43816 and *rcsB* gene deletion strains (\*\*\*\**p*<0.0001, *t*-test)

### Drug sensitivity test results of *Klebsiella pneumoniae* ATCC43816 and Δ*rcsB*

The drug sensitivity test results showed that the MIC values of imipenem, ampicillin/sulbactam and tegacyclin in the *rcsB* gene deletion strain were lower than those in the wild strain (Table 3).

**Table 3.**
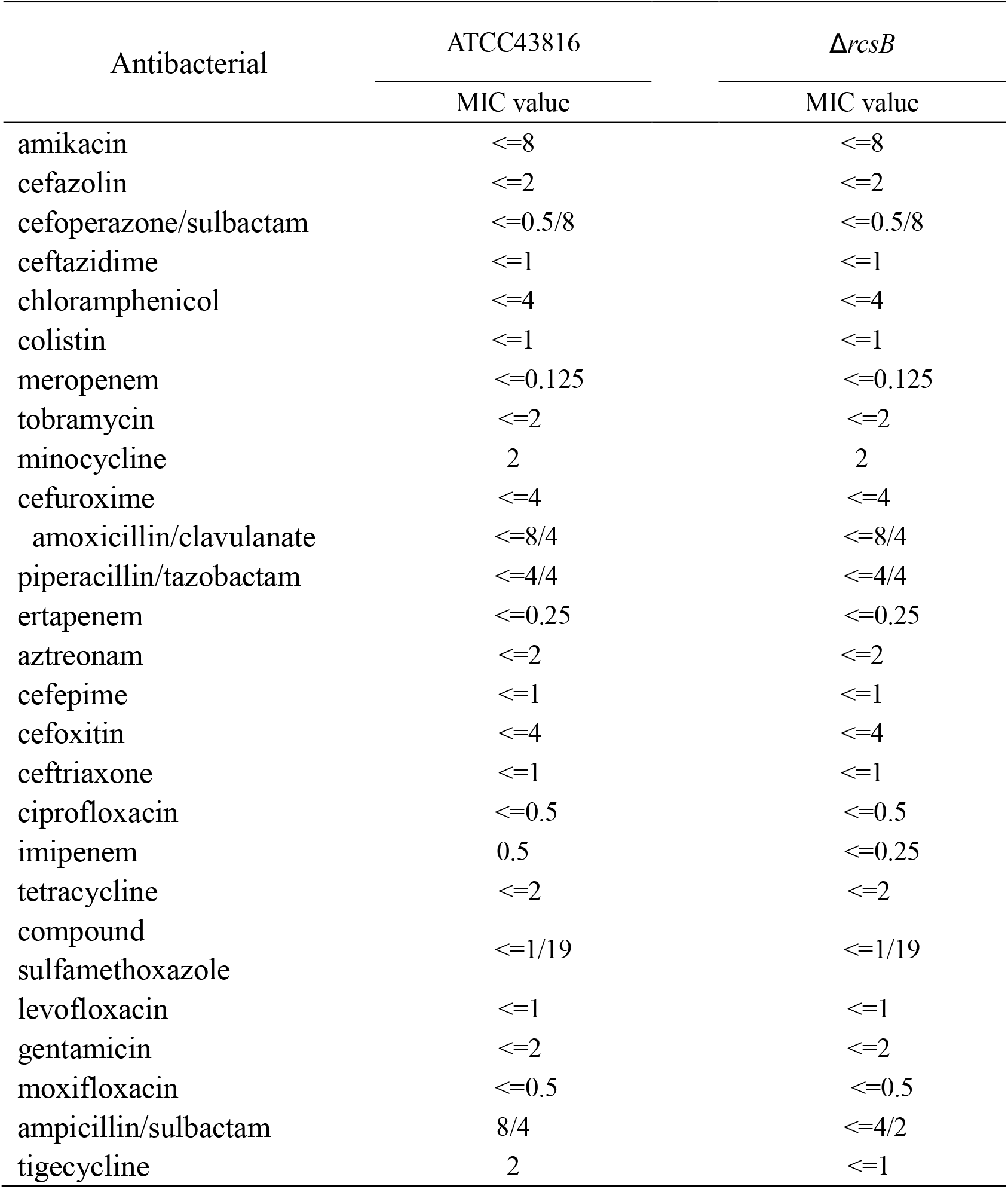
Comparison of drug sensitivity test results of *Klebsiella pneumoniae* ATCC43816 and Δ*rcsB*.

## Discussion

This study reviewed and analyzed 1,077 strains of pathogenic bacteria isolated from a hospital in Lianyungang in 2021, mainly gram-negative bacteria, focused on the drug sensitivity of *Klebsiella pneumoniae*, and found that its drug resistance rate to cefazolin was the highest. However, the detection rate of carbapenem resistant *Klebsiella pneumoniae* was as high as 40.0%, higher than the 37.9% in the statistics of Gaosaisai *et al*. It is considered that the reason may be that the intensive care unit (ICU) takes the first place in the sample source departments, and the number of samples was significantly higher than other departments. In this study, *Klebsiella pneumoniae* was taken as the experimental object, and the biological function of *rcsB* in highly virulent *Klebsiella pneumoniae* ATCC43816 was preliminarily explored by using growth curve, viscosity correlation test, uronic acid quantitative test and drug sensitivity test to compare and analyze the above biological characteristics of wild strains and deletion strains. It was found that the function of *rcsB* was not completely consistent with that reported in other studies. Lu Qin *et al*. found that the deletion of *rcsB* gene caused the growth rate of *Salmonella pullorum* to slow down, and the growth curve results in this study showed that the deletion of *rcsB* gene did not affect the growth rate of wild strains.

Andrea Rahn *et al*. found that *Escherichia coli K30* used *rcsB* to regulate capsule synthesis^[2]^. I Virlogeux *et al*. confirmed that *rcsB* was involved in the transcription of genes that activate capsule polysaccharide Vi polymer synthesis in *Salmonella typhimurium*^[3]^, which is consistent with the results of this study that *rcsB* gene deletion leads to reduced capsule synthesis in highly virulent *Klebsiella pneumoniae*. The change trend of high viscosity phenotype of *rcsB* gene deletion plants was consistent with the amount of capsule formation, suggesting that the high viscosity phenotype might be related to the amount of capsule formation. The drug sensitivity test results of *rcsB* showed that the increased sensitivity of the strain to some drugs after the deletion of *rcsB* gene may be due to the reduced synthesis of the capsule, which can help bacteria resist the bactericidal effect of antibacterial peptides and various inflammatory factors, phagocytosis of cells, and complement mediated bacteriolysis^[4-10]^. Therefore, the drug resistance of *Klebsiella pneumoniae* could be improved by inhibiting the expression of *rcsB* by some means.

To sum up, it is urgent to strengthen the rational use of antibiotics to improve the drug resistance of *Klebsiella pneumoniae* in this hospital, and the improvement of drug sensitivity of strains after *rcsB* gene deletion may provide new ideas.

### Data analysis

All numerical data were plotted as mean ± SEM unless otherwise indicated. The statistical analyses were performed using GraphPad Prism 8. Determination of significance between groups was performed using Student *t*-tests as indicated.

## Author Contributions

HQH drafted the manuscript. TW, JZ, SML and HD revised the manuscript critically for important intellectual content. All authors contributed to the article and approved the submitted version.

